# Impact of implementation choices on quantitative predictions of cell-based computational models

**DOI:** 10.1101/092924

**Authors:** Jochen Kursawe, Ruth E. Baker, Alexander G. Fletcher

## Abstract

‘Cell-based’ models provide a powerful computational tool for studying the mechanisms underlying the growth and dynamics of biological tissues in health and disease. An increasing amount of quantitative data with cellular resolution has paved the way for the quantitative parameterisation and validation of such models. However, the numerical implementation of cell-based models remains challenging, and little work has been done to understand to what extent implementation choices may influence model predictions. Here, we consider the numerical implementation of a popular class of cell-based models called vertex models, which are often used to study epithelial tissues. In two-dimensional vertex models, a tissue is approximated as a tessellation of polygons and the vertices of these polygons move due to mechanical forces originating from the cells. Such models have been used extensively to study the mechanical regulation of tissue topology in the literature. Here, we analyse how the model predictions may be affected by numerical parameters, such as the size of the time step, and non-physical model parameters, such as length thresholds for cell rearrangement. We find that vertex positions and summary statistics are sensitive to several of these implementation parameters. For example, the predicted tissue size decreases with decreasing cell cycle durations, and cell rearrangement may be suppressed by large time steps. These findings are counter-intuitive and illustrate that model predictions need to be thoroughly analysed and implementation details carefully considered when applying cell-based computational models in a quantitative setting.

## 1 Introduction

Computational modelling is increasingly used in conjunction with experimental studies to understand the self-organisation of biological tissues [1, 2]. Popular computational models include ‘cell-based’ models that simulate tissue behaviour with cellular resolution. Such models naturally capture stochastic effects and heterogeneity when only few cells are present and can be used to explore tissue behaviour when complex assumptions on the cellular scale prevent us from deriving continuum approximations on the tissue scale. The applications of cell-based models range from embryonic development [3–7], to wound healing [8] and tumour growth [9]. However, the numerical solution of cell-based models remains challenging since multi-scale implementations of such models, coupling processes at the subcellular, cellular, and tissue scales, may suffer from numerical instabilities [10, 11], and many such models include parameters of numerical approximation or parameters that have no direct physical correlate. These issues are of growing importance as cell-based models become used in an increasingly quantitative way [12–14]. Thus, we need to be aware of any impacts that numerical implementation choices may have on model predictions.

Here, we analyse a well-established class of cell-based model, the vertex model [15], to understand to what extent choices of numerical implementation and non-physical model parameters may affect model predictions. Vertex models were originally developed to study inorganic structures, such as foams [16] and grain boundaries [17,18], where surface tension and pressure drive dynamics. They have since been modified to study epithelial tissues [19–22], one of the major tissue types in animals. Epithelia form polarized sheets of cells with distinct apical (‘top’) and basal (‘bottom’) surfaces, with tight lateral attachments nearer their apical surface. The growth and dynamics of such sheets play a central role in morphogenesis and wound healing, as well as in disease; for example, over 80% of cancers originate in epithelia [23]. In two-dimensional vertex models, epithelial cell sheets are approximated by tessellations of polygons representing cell apical surfaces, and vertices (where three or more cells meet) move in response to forces due to growth, interfacial tension and hydrostatic pressure within each cell (figure 1A-C). Vertex models typically include cell growth and proliferation. In addition, cells exchange neighbours through so-called T1 transitions (figure 1D) whenever the length of a cell-cell interface falls below a threshold, and any triangular cell whose area falls below a threshold is removed by a so-called T2 transition (figure 1E).

**Figure 1:**
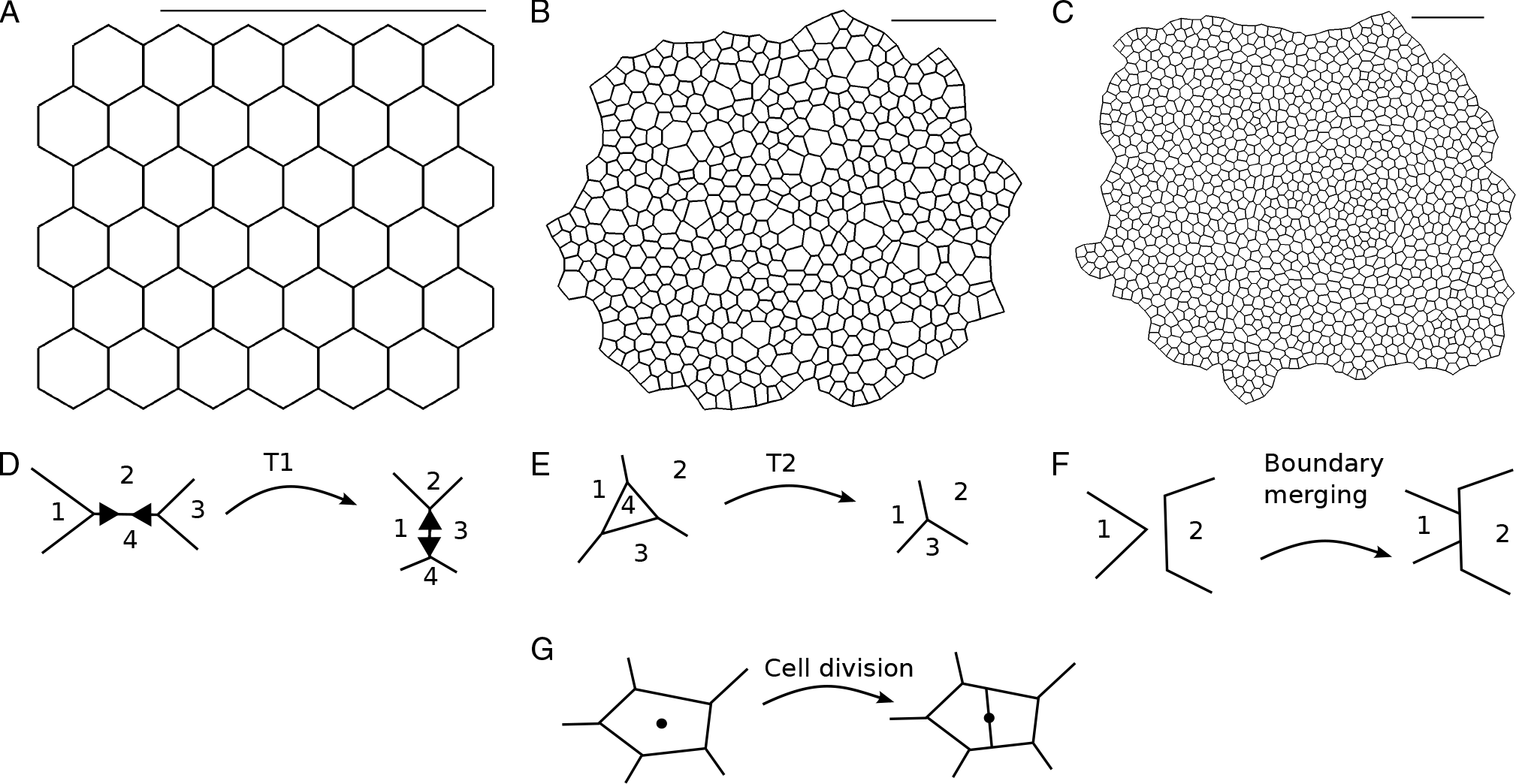
Two-dimensional vertex models represent cells in an epithelial tissue as polygons and allow different types of vertex rearrangement. (A-C) Snapshots of an example vertex model simulation used in our analysis. The growing *in silico* tissue undergoes five rounds of cell division. (A) The initial condition is a hexagonal packing of 36 cells. (B) Simulation progress after 6,750 time units at an intermediate stage of tissue growth. The tissue boundary is allowed to move freely and individual cells grow before division. (C) Snapshot of the tissue at the end of the simulation at 27,000 time units. After the fifth (last) round of divisions the tissue relaxes into a stable configuration. Simulated tissues in (B-C) are rescaled to fit the view, a scale bar of fixed length is added for comparison. Parameter values are listed in table 1. Throughout the simulation, vertices may rearrange by T1 transitions (D), T2 transitions (E), boundary merging (F), and cell division (G).

Vertex models have been used to study a variety of processes in epithelial tissues [3–6,24–38]. These processes include growth of the *Drosophila* wing imaginal disc [3, 4], migration of the visceral endoderm of mouse embryos [5], and tissue size control in the *Drosophila* embryonic epidermis [31]. A common approach in such studies is to consider forces on vertices arising as a result of minimizing the total stored energy in the tissue. The functional form for this total stored energy varies between applications, but is typically chosen to reflect the effect of the force-generating molecules which localise at or near the apical surface. This energy function is then used either to derive forces that feed into a deterministic equation of motion for each vertex, which must be integrated over time [4, 24, 28], or else minimized directly assuming the tissue to be in quasistatic mechanical equilibrium at all times [3, 25]. A third approach is to apply Monte Carlo algorithms to find energy minima [39, 40].

Previous theoretical analyses of vertex models have elucidated ground state configurations and their dependence on the mechanical parameters of the model [41], inferred bulk material properties [42–44], and introduced ways to superimpose finite-element schemes for diffusing signals with the model geometry [45]. In other work, vertex models have been compared to lattice-based cellular Potts models and other cell-based modelling frameworks [46, 47].

In the case of vertex models of grain boundaries, the authors of [18] proposed an adaptive time-stepping algorithm to accurately resolve vertex rearrangements without the need of adhoc rearrangement thresholds and provide a numerical analysis of the simulation algorithm. However, vertex models in that context only consider energy terms that are linear in each grain-grain (or cell-cell) interface length, whereas the energy terms in vertex models of biological cells typically depend non-linearly on cell areas and perimeters.

Importantly, previous studies such as [18] do not analyse to what extent changes in hidden model parameters, such as parameters of numerical approximation, like the size of the time step, or non-physical model parameters, such as length thresholds for cell rearrangement, can influence vertex configurations and other summary statistics. Here, we analyse a force-propagation implementation of vertex models [48, 49] as applied to a widely studied system in developmental biology, the larval wing disc of the fruit fly *Drosophila* [3, 4, 25]. We conduct convergence analyses of vertex positions with respect to all numerical and non-physical model parameters, and further analyse to what extent experimentally measurable summary statistics of tissue morphology, such as distributions of cell neighbour numbers and areas, depend on these parameters.

We find that vertex model predictions are sensitive to the length of cell cycle duration, the time step, and the size of the edge length threshold for cell rearrangement. Specifically, vertex configurations do not converge as the time step, the edge length threshold for cell rearrangement, or the area threshold for cell removal are reduced. For example, reductions in the cell cycle duration may promote cell removal and reduce the size of the simulated tissue by up to a factor of two. We find that both the size of the time step and the size of the edge length threshold can influence the rate of cell rearrangement. Counterintuitively, the rate of cell removal is robust to changes in the area threshold for cell removal over multiple orders of magnitude. Further, analysing the active forces within the tissue reveals that vertices are subject to stronger forces during periods when cells grow and divide.

The remainder of the paper is organised as follows. In section 2, we describe our vertex model implementation of growth in the *Drosophila* larval wing disc. In section 3 we present our results. Finally, we discuss our results and draw conclusions for the use of cell-based models in quantitative biology in sections 4 and 5.

## 2 Methods

We consider a vertex model of the growing *Drosophila* wing imaginal disc, a monolayered epithelial tissue that is one of the most widely used applications of vertex models. The wing imaginal disc initially comprises around 30 cells, and undergoes a period of intense proliferation until there are around 10,000 or more cells [3, 25]. Here, we outline the technical details of our model implementation. We start by introducing the equations of motion, then describe the initial and boundary conditions and implementations of cell growth and neighbour exchange.

### Equations of motion

In two-dimensional vertex models epithelial tissues are represented as tessellations of polygons that approximate the apical cell surfaces. We propagate the position of each vertex over time using an overdamped force equation, reflecting that cell junctions are not associated with a momentum. The force equation takes the form

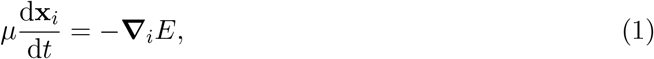
where *μ* is the friction strength, **x***_i_*(*t*) is the position vector of vertex *i* at time *t*, and *E* denotes the total stored energy. The number of vertices in the system may change over time due to cell division and removal. The symbol **∇***_i_* denotes the gradient operator with respect to the coordinates of vertex *i*. The total stored energy takes the form 

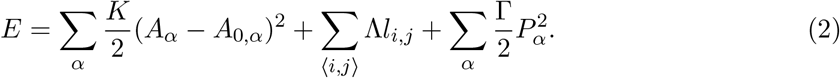

Here, the first sum runs over every cell *α* in the tissue, *A_α_* denotes the area of cell *α* and *A*_0,*α*_ is its target area. This term penalises deviations from the target area for individual cells, thus describing cellular bulk elasticity. The second sum runs over all cell edges 〈*i, j*〉 in the sheet and penalizes long edges (we choose ˄ *>* 0), representing the combined effect of binding energy and contractile molecules at the interface between two cells. The third sum also runs over all cells, and *P_α_* denotes the perimeter of cell *α*. This term represents a contractile acto-myosin cable along the perimeter of each cell [3]. The parameters *K*, Λ, and Γ together govern the strength of the individual energy contributions.

Before solving the model numerically, we non-dimensionalise it to reduce the number of free parameters [3]. Rescaling space by a characteristic length scale, *L*, chosen to be the typical length of an individual cell, and time by the characteristic timescale, *T* = *μ/KL*^2^, equations (1) and (2) become

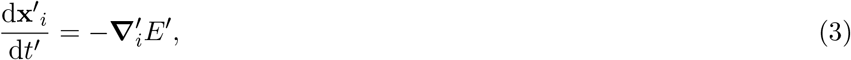

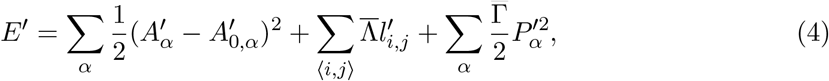

where 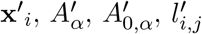 and *P′_α_* denote the rescaled *i*^th^ vertex positions, the rescaled area and target area of cell *α*, the rescaled length of edge 〈*i, j*〉, and the rescaled cell perimeter of cell *α*, respectively. The symbol 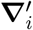 denotes the gradient with respect to the rescaled *i*^th^ vertex position. In the non-dimensionalised model, cell shapes are governed by the rescaled target area of each cell *A′*_0,*α*_ and the rescaled mechanical parameters, 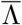 and 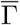. For these parameters we use previously proposed values [3], unless stated otherwise. A complete list of parameters used in this study is provided in table 1.

To solve equations (3) and (4) numerically we use a forward Euler scheme:

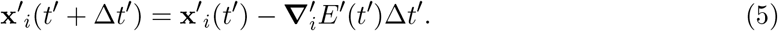

We analyse the dependence of simulation outcomes on the size of ∆*t′* in the Results section.

### Initial and boundary conditions

Initially, the sheet is represented by a regular hexagonal lattice of six by six cells (figure 1A). The boundary of the lattice is allowed to move freely throughout the simulation. Each cell has initial area and target area 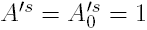, respectively.

### Cell neighbour exchange and removal

T1 transitions (figure 1D) are executed whenever the length of a given edge decreases below the threshold *l′*_T1_ = 0.01. The length of the new edge, *l*_new_ = *ρ l*_T1_ (*ρ* = 1.5), is chosen to be slightly longer than this threshold to avoid an immediate reversion of the transition.

A second topological rearrangement in vertex models is a T2 transition, during which a small triangular cell or void is removed from the tissue and replaced by a new vertex (figure 1E). In our implementation any triangular cell is removed if its area drops below the threshold *A′*_T2_ = 0.001. The energy function, equation (2), in conjunction with T2 transitions can be understood as a model for cell removal: cells are extruded from the sheet by a T2 transition if the energy function, equation (2), leads to a sufficiently small cell. Note that in equation (2) the bulk elasticity or area contribution of a cell *α* is finite even when the area *A_α_* is zero, allowing individual cells to become arbitrarily small if this is energetically favourable. As cells decrease in area they typically also reduce their number of sides. Hence, it is sufficient to remove only small triangular cells instead of cells with four or more sides [3, 4, 25].

We further model the merging of overlapping tissue boundaries (figure 1F). Whenever two boundary cells overlap, a new edge of length *l*_new_ is created that is shared by the overlapping cells. In cases where the cells overlap by multiple vertices, or if the same cells overlap again after a previous merging of edges, the implementation ensures that two adjacent polygons never share more than one edge by removing obsolete vertices. The merging of boundary edges is discussed in further detail in [48].

### Cell growth and division

Unless stated otherwise the tissue is simulated for *n*_*d*_ = 5 rounds of division, i.e. each cell divides exactly *n*_*d*_ times. To facilitate comparison with previous simulations of the wing disc where vertices were propagated by minimising the energy function (2) [3, 41], we model each cell to have two cell cycle phases: quiescent and growing. The duration of the first, quiescent, phase of the cell cycle is drawn independently from an exponential distribution with mean 2*t′_l_/*3, where *t′_1_* is the total cell cycle duration. We introduce stochasticity in this phase of the cell cycle to avoid biologically unrealistic synchronous adjacent divisions; this also helps keeping the simulations in a quasistatic regime since adjacent divisions are prevented from influencing each other, thus maintaining mechanical equilibrium. The duration of the second, growing, phase of the cell cycle is fixed at length *t′_l_/*3 for each cell. During this time the target area, *A′*_0,*α*_, of the cell grows linearly to twice its original value.

Upon completion of the growth phase, the cell divides. We choose a fixed duration for the growth phase to ensure gradual, quasistatic cell growth. Two-stage cell cycles with an exponentially distributed and a fixed length contribution have previously been observed in various cell cultures [50, 51] and have been applied to model growth in the *Drosophila* wing imaginal disc [28].

The assigning of these cell cycle stages to two thirds and one third of the total cell cycle duration *t′*_l_, respectively, allows us to modify the average age of a dividing cell with a single parameter. This decomposition of the cell cycle ensures that cell cycle durations are stochastic, while allowing the growth phase to occupy a significant proportion of the total cell cycle duration, ensuring gradual, quasistatic growth. The assumption that the tissue is in a quasi-steady state is common in vertex models [3, 27, 28, 34] and reflects the fact that the time scales associated with mechanical rearrangements (seconds to minutes) are an order of magnitude smaller than typical cell cycle times (hours) [3].

At each cell division event, a new edge is created that separates the newly created daughter cells (figure 1G). The new edge is drawn along the short axis of the polygon that represents the mother cell [48]. The short axis has been shown to approximate the division direction (cleavage plane) of cells in a variety of tissues [52], including the *Drosophila* wing imaginal disc [53]. The short axis of a polygon crosses the centre of mass of the polygon, and it is defined as the axis around which the moment of inertia of the polygon is maximised. Each daughter cell receives half the target area of the mother cell upon division.

Applying this cell cycle model, we let the tissue grow for *n*_*d*_ = 5 generations until it contains approximately 1,000 cells, making it sufficiently large to obtain summary statistics of cell packing. Note that the precise number of cells at the end of the simulation varies, due to variations in the number of T2 transitions by which individual cells are removed from the tissue. Each cell of the last generation remains in the quiescent phase of the cell cycle until the simulation stops. We select the total simulation time to be *t′*_tot_ = 27, 000, unless specified otherwise. This duration is chosen such that the tissue can relax into its equilibrium configuration after the final cell division.

## Computational implementation

We implement the model within Chaste, an open source C++ library that provides a systematic framework for the simulation of vertex models [48, 49]. Our code is available in the supplementary material as a zip archive. Pseudocode for our implementation is provided in algorithm 1. Each time step starts by updating the cell target areas. Then, cell division, removal (T2 transitions), rearrangement (T1 transitions), and boundary merging are performed before incrementing the simulation time. The algorithm stops when the end time of the simulation is reached.

**Algorithm 1:**
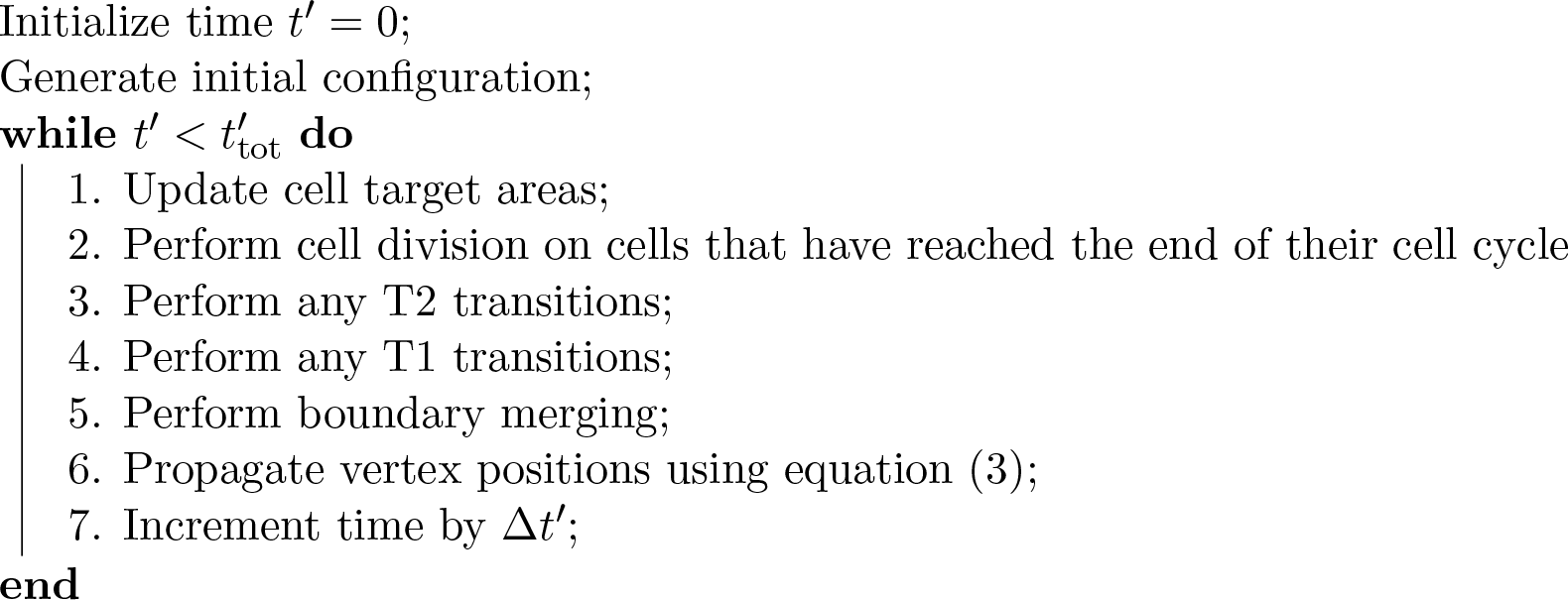
Pseudocode of the simulation algorithm.

**Table 1.** Description of parameter values used in our simulations.

| Parameter | Description | Value | Reference |
| --- | --- | --- | --- |
| $\bar{\Lambda}$ | Cell-cell adhesion coefficient | 0.12 | [3] |
| $\bar{\Gamma}$ | Cortical contractility coefficient | 0.04 | [3] |
| $\Delta t'$ | Time step | 0.01 | [48] |
| $A'_{\min}$ | T2 transition area threshold | 0.001 | [48] |
| $l'_{\text{T1}}$ | T1 transition length threshold | 0.01 | [48] |
| $\rho$ | New edges after a T1 transition have the length $l'_{\text{new}} = \rho l'_{\text{T1}}$ | 1.5 | [48] |
| $A'^s$ | Initial cell area | 1.0 | [3] |
| $A_0'^s$ | Initial cell target area | 1.0 | [3] |
| $N^s$ | Initial cell number | 36 | [3] |
| $t'_l$ | Mean cell cycle duration | 1,750 | – |
| $t'_{\text{tot}}$ | Simulation duration | 27,000 | – |
| $n_d$ | Total number of divisions per cell | 4 | – |

For parameter values for which no reference is given, please see main text for details on how these values were estimated. Spatial and temporal parameters are non-dimensionalised (see section 2 for details).

## 3 Results

In this section, we analyse how model behaviour depends on numerical and non-physical model parameters. Vertex models are typically used to predict summary statistics of cell packing and growth, such as the distribution of cell neighbour numbers and areas [3, 25]. We analyse how these summary statistics depend on simulation parameters. Specifically, we focus on the final number of cells in the tissue, the total tissue area, the numbers of cell rearrangements (T1 transitions) and cell removals (T2 transitions), the distribution of cell neighbour numbers, and the correlation between cell neighbour number and cell area. Note that we exclude cells on the tissue boundary from statistics of cell neighbour numbers in order to avoid boundary artefacts, which can be seen in figure 1C. In figure 1C, cell shapes along the tissue boundary differ from those in the bulk of the tissue, and the cell neighbour number is poorly defined for cells along the tissue boundary, since it does not coincide with the number of cell edges.

### Tissue size is sensitive to cell cycle duration

In previous vertex model applications [3, 4, 25], experimentally measured summary statistics of cell packing were reproduced using an energy minimisation implementation. Such energy minimisation schemes assume quasistatic evolution of the sheet, where the tissue is in mechanical equilibrium at all times. It is unclear to what extent summary statistics are preserved when the tissue evolves in a dynamic regime.

We analyse the dependence of the summary statistics on the cell cycle duration, *tl*, in figure 2. The cell number and tissue area at the end of the simulation, and the total number of cell rearrangements, vary by up to a factor of two as the mean cell cycle duration increases from five to 2000 non-dimensional time units (figure 2A-D). The cell number and tissue area increase with the mean cell cycle duration, whereas the amount of rearrangement (T1 transitions) decreases, reflecting a reduction in cell removal events (T2 transitions). The cell number and the tissue area do not increase further for mean non-dimensional cell cycle durations larger than 1,000 time units. In this regime, the total number of rearrangements and cell removals also cease decreasing. We thus identify this regime as the quasistatic regime, where the tissue maintains mechanical equilibrium throughout the simulation. Note, however, that neither the total cell number, nor the tissue area, the number of cell rearrangements or the number of cell removal converge numerically as the mean cell cycle duration increases, due to the stochastic nature of the system.

**Figure 2:**
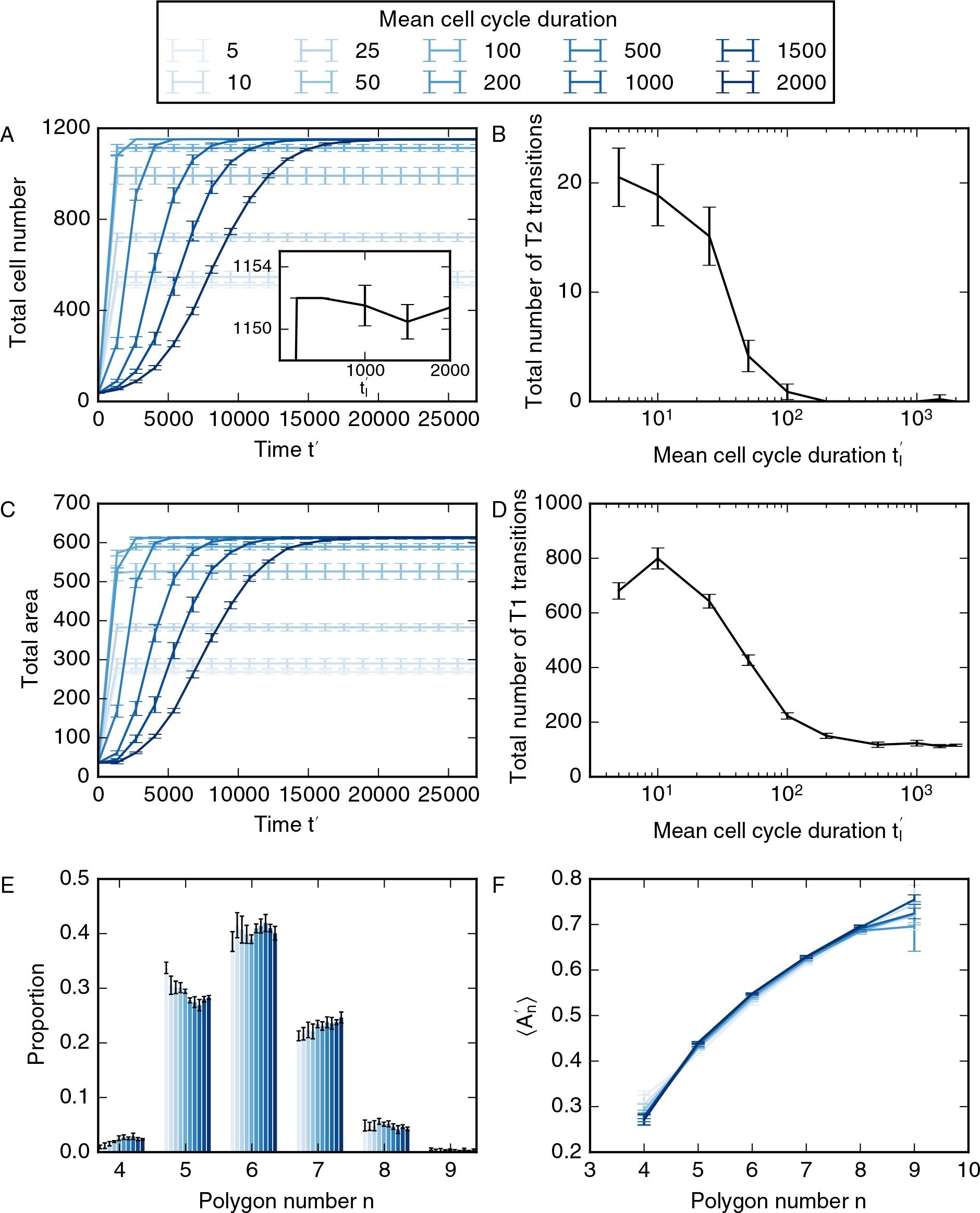
Variation of cell numbers (A), number of T2 transitions (B), tissue area (C), total number of T1 transitions (D), cell neighbour number distribution (E) and mean area per polygon class (F) with mean cell cycle duration. Error bars denote standard deviations across 100 simulations. All simulation parameters are provided in table 1.

The cell neighbour number distribution depends on the cell cycle duration in a non-linear fashion (figure 2E). For example, the number of hexagons peaks at cell cycle durations of 10 as well as 1,000 time units. For cell cycle durations longer than 1,000 time units the numbers of pentagons and heptagons increase as the cell cycle duration increases, while the number of hexagons decreases. We interpret this non-linear dependence as resulting from changes in cell neighbour numbers due to cell division and due to cell neighbour exchanges. As the cell cycle duration exceeds *t′*_l_ = 10, a decrease in the number of cell removal events leads to an increase in cell division events which, in turn, drives the polygon distribution away from its hexagonal initial condition. As the number of cell divisions ceases to increase the number of cell rearrangements drops as well, and the number of hexagons reaches a second peak. Increasing the time between cell divisions further decreases the number of hexagons. Note that none of the simulated polygon histograms coincide with previously reported histograms in which pentagons outweigh hexagons [3, 25], despite choosing identical parameters in energy equation (2). We discuss possible reasons for this difference in section 4.

Another common summary statistic of cell packing is the mean area of cells of each polygon number 〈*A′*_n_〉, where 〈*·*〉 denotes an average across all cells in the tissue that are not on the tissue boundary, *A′* is the rescaled cell area, and *n* is the polygon number, i.e. the number of neighbours that each cell has. This summary statistic is often used to characterise epithelia [3, 26, 54, 55]. We find that the mean cell area for each polygon number is not sensitive to changes in cell cycle length and increases monotonically with polygon number (figure 2F).

We interpret the data in figure 2 as follows. Differences in tissue size and cell packing arise due to a sensitive interplay between the cell cycle duration and the timescale for mechanical relaxation of the tissue, *T*. Growing cells push against their neighbours, leading to tissue growth. This outward movement is counteracted by the friction term in the force equation (1). As cells grow more quickly, i.e. with smaller cell cycle durations, the force required to push the surrounding cells outward increases. For sufficiently small cell cycle durations, the forces may become strong enough to cause cell extrusion. This finding is may not be biologically relevant when studying growth in the *Drosophila* wing imaginal disc, since in this system the time scales for mechanical rearrangement are orders of magnitude smaller than the time scales associated with growth and proliferation [3]. However our results suggest that, in other systems, where cells divide on the time scales of minutes rather than hours, such as the *Drosophila* embryonic epidermis, cell extrusion may be induced during periods of fast tissue growth.

### Cell growth and division increase forces within the tissue

The energy expression (4) leads to three different force contributions on each vertex: an area force; an edge force; and a perimeter force. In figure 3 we analyse the magnitude of these contributions for a simulation with mean cell cycle duration *t′_l_* = 2000. The solid line represents the average magnitudes for the individual contributions for all forces in the tissue, and the shaded areas mark one standard deviation. The strongest force contribution is the area force (figure 3A), whereas the weakest is the edge force (figure 3B). This relationship is intuitive if one considers the directions of the individual force contributions when both 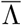 and 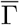 are positive: Most cells in the tissue have areas smaller than their target area of 1.0 (compare with figure 2F), hence for an individual cell, the area force contribution points outwards from the cell. The edge contribution and perimeter contribution (figure 3C) point inwards for individual cells, thus counteracting the area force. It follows that the area contribution is strongest since, in mechanical equilibrium, it counteracts the sum of the edge and perimeter contributions. The variation of each force contribution has the same order of magnitude as their mean values, illustrating that the forces on vertices can vary strongly across the tissue. The force magnitudes change throughout the simulation, and they peak at a value that is 50% higher than the final values. For times larger than 15000 time units, the forces do not change with time in figure 3. At this time cells stop dividing and the final cell number is reached, illustrating that the forces are largest when the tissue size is increasing most rapidly. This transient rise in forces emerges because cells in the interior of the simulated tissue push on their neighbours as they grow before division. These observations enable us to predict that cells undergoing active processes, such as growth and division, are subject to significantly higher forces than cells in quiescent tissues.

**Figure 3:**
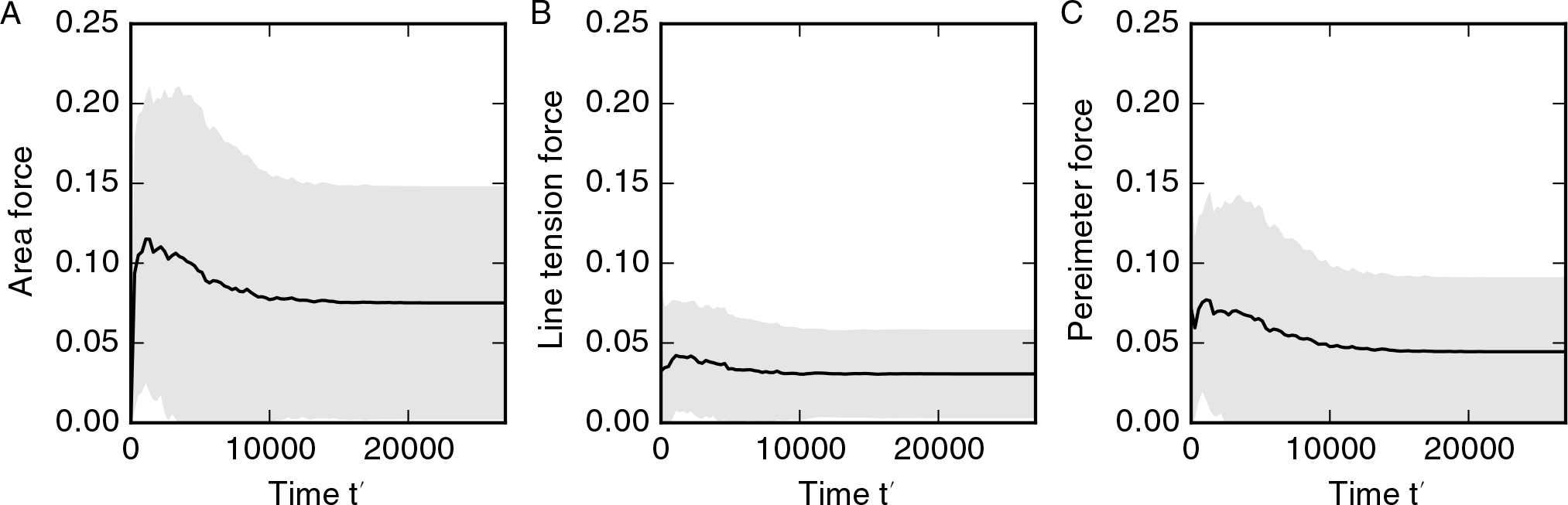
Magnitude of area (A), edge (B), and perimeter force (C) contributions over time. The solid lines represent the average of force contribution magnitudes across all vertices of one simulation. The shaded regions represent one standard deviation of the force contribution magnitudes across the tissue. A cell cycle duration of *t′*_l_ = 2000 is used. All other parameters are listed in table 1.

### Large time steps suppress cell rearrangement

When using an explicit Euler method to propagate the model forward in time, such as in equation (5), the time step should be chosen sufficiently small to provide a stable and accurate numerical approximation of the model dynamics. To this end, we conduct a convergence analysis. To reduce simulation times, we conduct the convergence analysis on sample simulations in which each cell divides *n*_*d*_ = 4 times instead of five, and set the total simulation time as *t′*_tot_ = 21, 000. We choose a series of decreasing time steps, ∆*t′_k_*, and define the error function

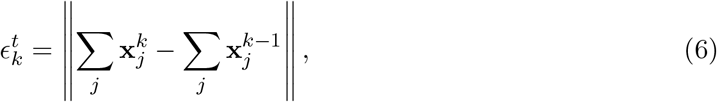

where the sums run over all vertex positions, 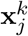, at the end of the simulation with time steps ∆*t′_k_* and ∆*t′*_*k*-1_. The error function (6) evaluates the differences between the sums of final vertex positions at decreasing values of the time step. To ensure that simulations with consecutive values of the time step follow identical dynamics we generate fixed series of exponentially distributed random variates from which we calculate the cell cycle durations.

We plot results of our analysis of the convergence of the vertex positions with the time step ∆*t′* in figure 4. In general, the error function does not converge. However, for most simulations the error function (6) assumes values smaller than 10^−1^ for time steps smaller than 10^−2^ (figure 4A). Note that this time step is five orders of magnitude smaller than the average cell cycle duration. When the time step is larger than 10^−2^ the error function (6) is larger than one since a significant number of T1 transitions are suppressed. On rare occasions, for less than five examples out of 100, the error function may be non-negligible even if the time step is smaller than 10^−2^. These large values of the error function (6) reflect changes in the number of T1 transitions as the time step decreases (figure 4B). When the time step is smaller than 10^−2^ summary statistics of cell packing, such as the distribution of cell neighbour numbers (figure 4C) or the total number of cells, do not change as the time step is decreased further. Note that the distribution of cell neighbour numbers in figure 4C differs from those in figure 2 due to the decreased number of divisions per cell, *n*_*d*_. Further, we conclude from our analysis in figure 4 that it is necessary to use a time step smaller than 0.01 in order to arrive at physically meaningful solutions of the vertex model, since otherwise the amount of cell rearrangement and summary statistics of cell packing will be affected by the numerical implementation of the model.

**Figure 4:**
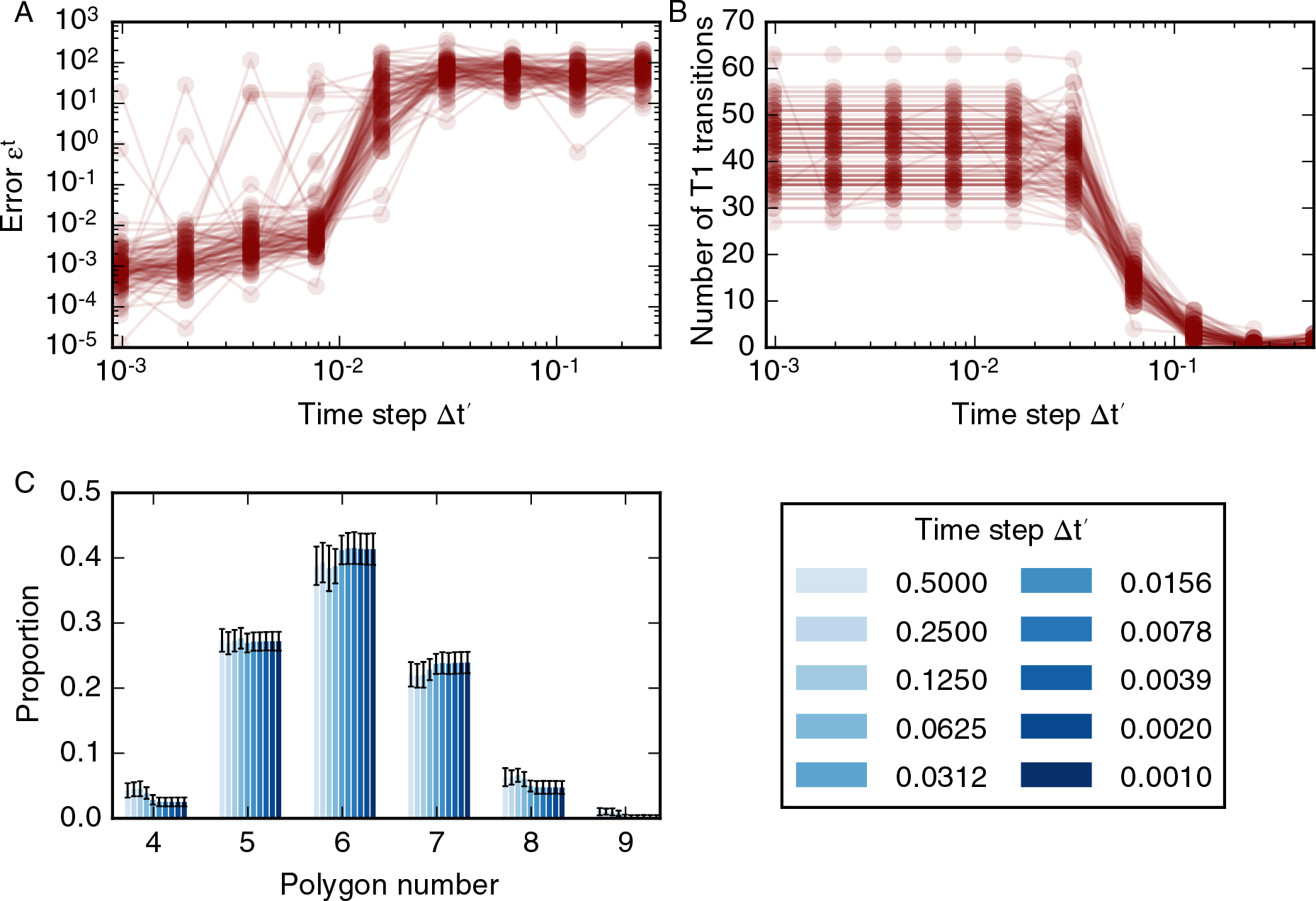
Variation in simulation result with the time step. (A) The error function (6) for 100 different realisations of the model plotted as overlapping, opaque curves. The error function decreases as the time step is decreased, but does not converge for all simulations. (B) The dependence of the number of T1 transitions on the time step for 100 model realisations. The number of T1 transitions in the simulations is stable for time steps smaller than 0.02 and decreases with time steps greater than 0.002. (C) For time steps ∆*t′* < 0.02 the cell neighbour number distribution is stable; the means of individual polygon class proportions vary by less than 0.01. In these simulations, cells undergo *n*_*d*_ = 4 rounds of division, and the total simulation time *t′*_tot_ = 21, 000. All other parameter values are listed in table 1. Error bars denote standard deviations across 100 simulations.

An example of how differences in the number of T1 transitions and final vertex positions can emerge when the time step is smaller than 0.01 is shown in figure 5. In this figure, a cell division occurs in two simulations using a time step of 0.004 (figure 5A) and a time step of 0.002 (figure 5B). Both simulations use the same, fixed, series of cell cycle times, and vertex positions in both simulations are similar over time up until the illustrated division. Here, and throughout, cells divide along their short axis. In this example, the short axis of the cell intersects the cell boundary close to an existing vertex. Due to differences in the vertex positions of the cell, the new vertex is created on different cell-cell interfaces as the size of the time step varies. As the simulation progresses, these different vertex configurations propagate towards different final tissue configurations, leading to differences in the total number of T1 transitions and the error function. In figure 4, differences in final vertex positions are observed for all considered values of the time step. However, such differences in vertex positions do not propagate through to tissue-level summary statistics such as the distribution of cell neighbour numbers or areas.

**Figure 5:**
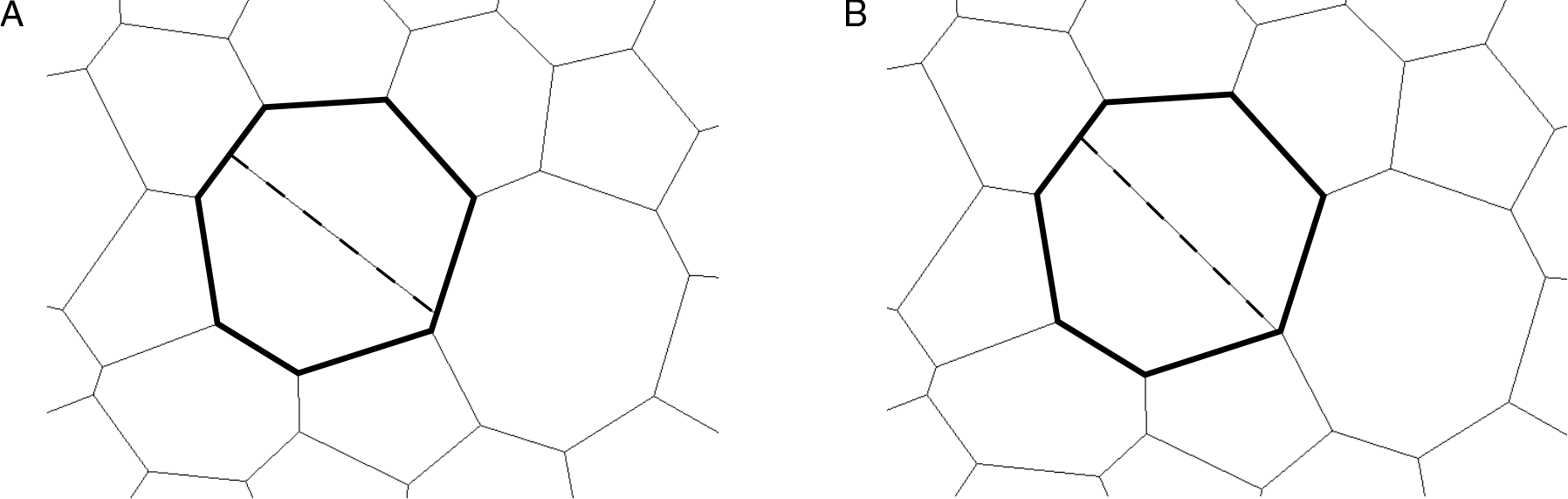
Differences in vertex configurations can arise in simulations run with different temporal resolution. A dividing cell in simulations run with time steps ∆*t′* = 0.004 (A) and ∆*t′* = 0.002 (B) is shown in bold. During the cell division, a new cell-cell interface (dashed line) is created along the short axis of the dividing cell by creating new vertices (see Methods section for details). The daughter cells of the dividing cell contain different vertices in the configurations corresponding to the two time steps. This leads to different vertex configurations at the end of the simulations.

### Model convergence with time step is not improved if higher-order numerical methods are used

The results in figures 4 and 5 were generated by propagating the vertex positions using a forward Euler time-stepping scheme. The choice of a forward Euler scheme over more accurate numerical methods is common in vertex models. For example, in a previous application where a tissue was relaxed starting from a random initial condition, it was shown that, in order to accurately resolve all T1 transitions, sufficiently small time steps had to be chosen that the benefits of higher order numerical methods were negligible [56]. However, in figures 4 and 5 vertex positions do not converge as the time step is decreased due to differences in T1 transitions and cell divisions for varying values of the time step, suggesting that convergence might be achieved if higher order numerical methods were used. We test this hypothesis in figure 6, where we record the error function (6) when propagating the vertex model with a fourth-order Runge-Kutta timestepping scheme as follows. First, all vertices are accumulated into the vertex vector **x***′*, such that if there are *N* vertices at time *t′* then the vector **x***′*(*t′*) has 2*N* components. We propagate the vertex vector using

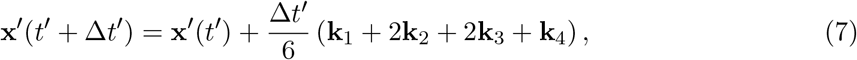

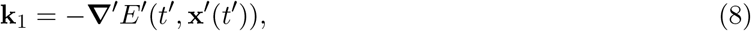

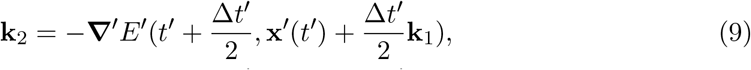

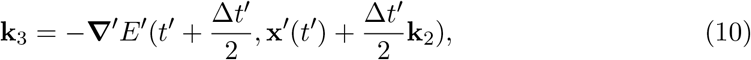

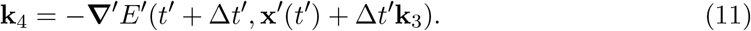

Here, **∇***′* denotes the gradient with respect to the vector **x**.

**Figure 6:**
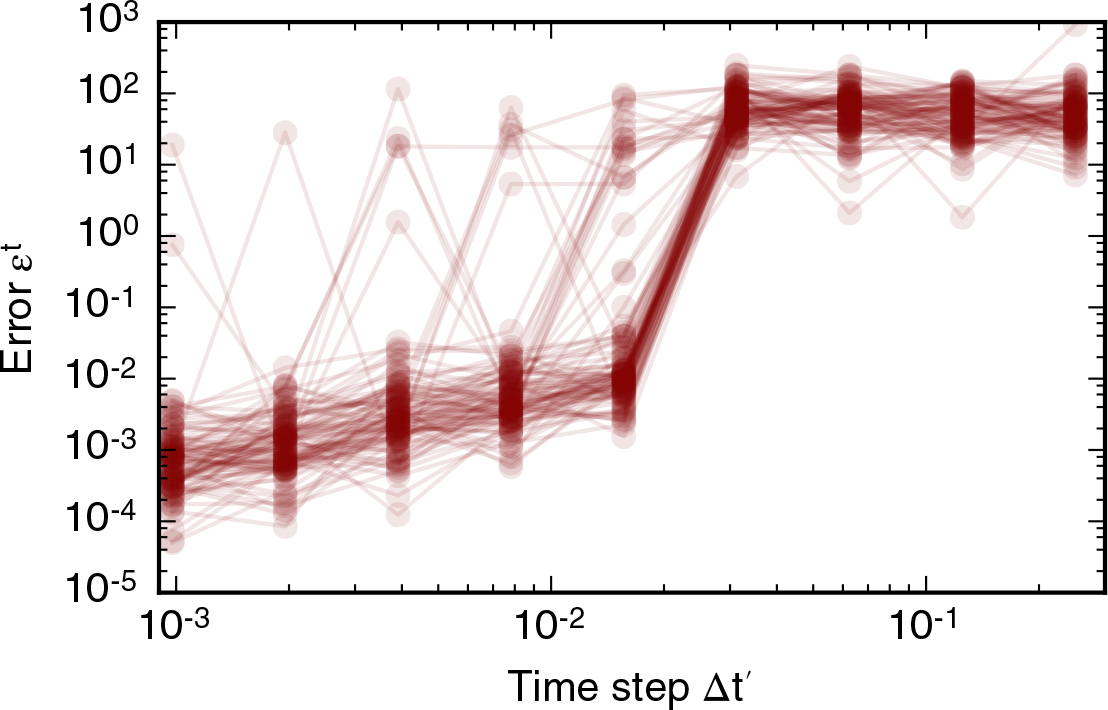
Variation in simulation result with the time step if a fourth-order Runge-Kutta scheme is used. The error function (6) for 100 different realisations of the model, evaluated using a fourth-order Runge-Kutta scheme, is plotted as overlapping, opaque curves. The error function decreases as the time step is decreased, but does not converge for all simulations. This result is similar for simulations run with a forward Euler scheme in figure 4A.

Similar to the error function obtained using a forward Euler numerical scheme in figure 4A, the error function obtained using a fourth-order Runge-Kutta numerical scheme in figure 6 assumes values smaller than one for time steps below 0.01, but does not converge as the time step is decreased further. Comparing figures 4A and 6 we conclude that a higher-order time-stepping scheme does not improve the accuracy of vertex model propagation, since both the forward Euler and the fourth-order Runge-Kutta scheme require time steps smaller than roughly 0.01 in order for the error function (6) to assume values smaller than one on average, while exhibiting a similar degree of variability across all simulations.

### Occurrence of cell rearrangements is regulated by rearrangement threshold

We further analyse the dependence of vertex positions and summary statistics on the T1 transition threshold, *l′*_T1_. Similar to the time step convergence analysis, we define a series of decreasing values of *l′*_T1,*k*_ and the error function 

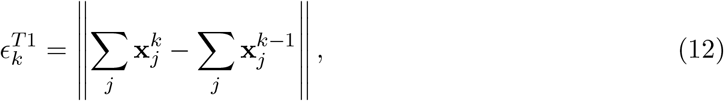

which measures the difference between the final vertex positions of simulations with decreasing values of the T1 transition threshold, *l′*_T1,*k*_. The variation of the error function with decreasing values of *l′*_T1,*k*_ is shown in figure 8A. For all considered values of *l′*_T1_ the error function does not converge and varies between values of 1 and 10^3^. Only for *l′*_T1,*k*_ < 10^−3^ is the error function (12) smaller than one for some simulations. However, for such small values of *l′*_T1_, many simulations fail as the simulation algorithm encounters situations that it cannot resolve, for example configurations including overlapping cells (figure 8B).

**Figure 7:**
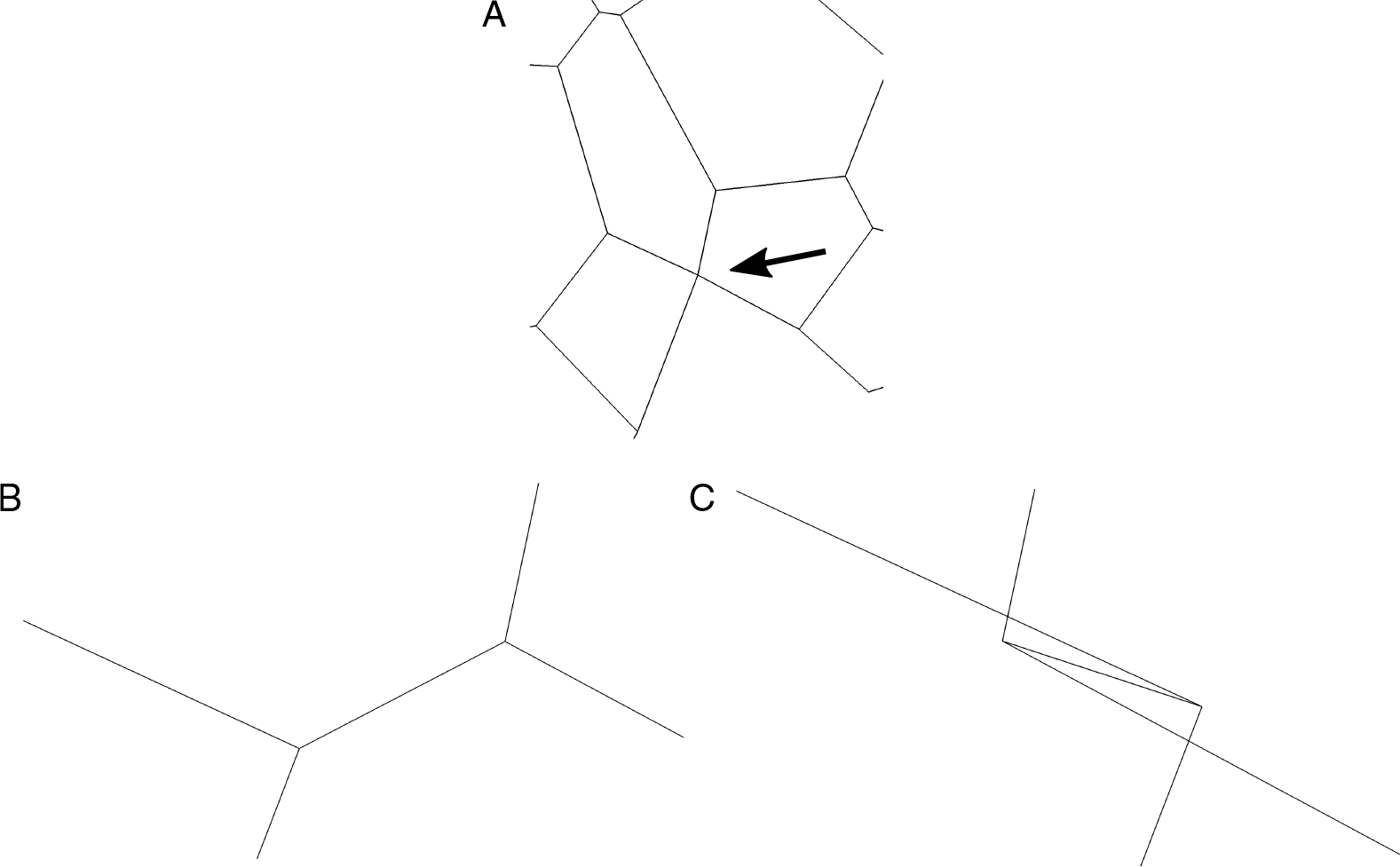
Small values of the T1 transition threshold, *l′*_T1_ *<* 10^−3^, suppress rearrangement and lead to failure of the simulation algorithm. One of the failing simulations in figure 8 is analysed. The tissue configuration in the last time step before simulation failure contains two vertices that appear to be merged due to a short edge on the tissue boundary. The short edge is indicated by an arrow (A) and magnified for the penultimate (B) and final completed time step (C) of the simulation. Since the short edge in the penultimate time step is prevented from rearranging, the two adjacent boundary cells intersect each other, leading to failure of the simulation.

**Figure 8:**
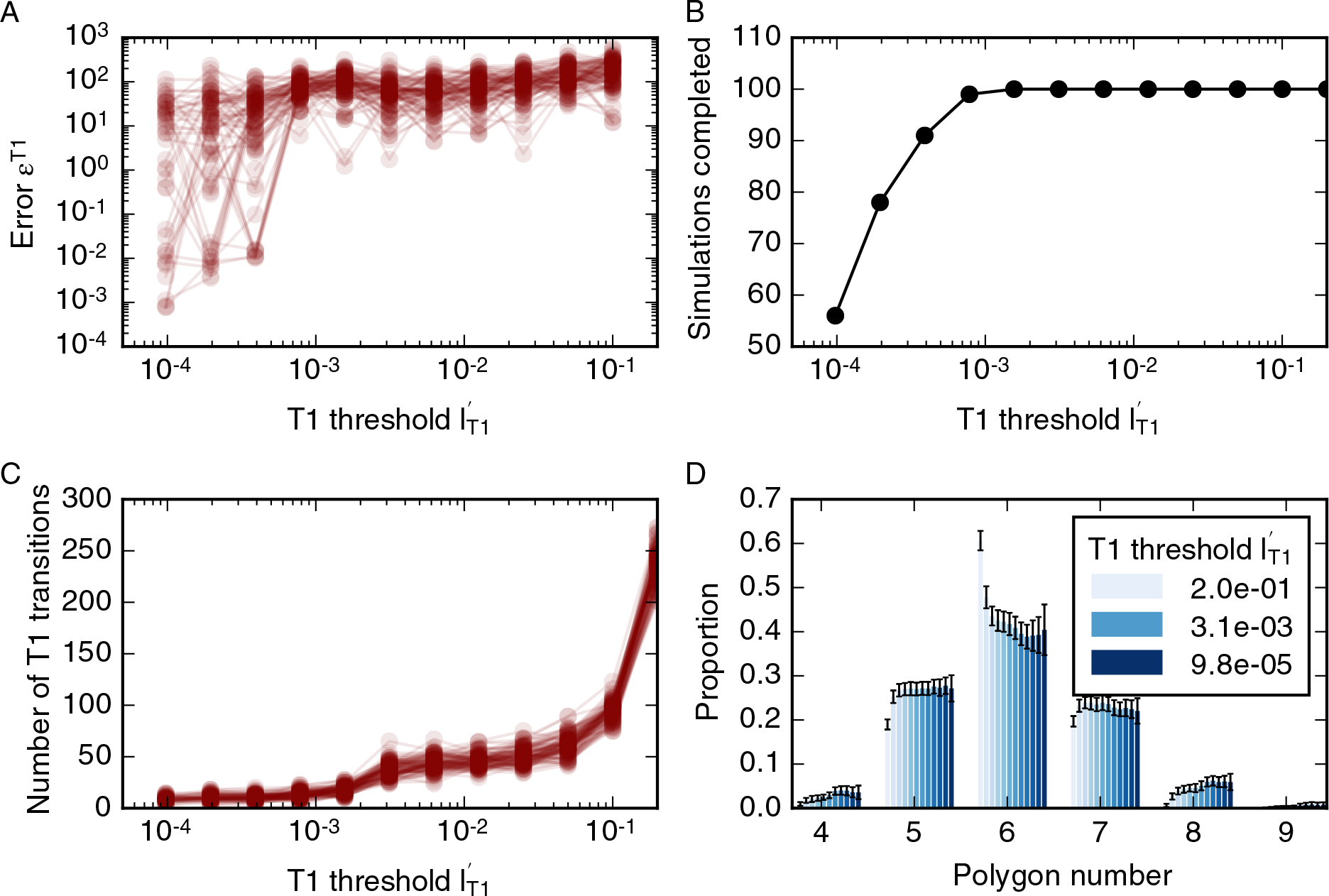
Variation of simulation result with size of the T1 transition threshold, *l′*_*T*1_. (A) The dependence of the error function on *l′*_*T*1_ for 100 model realisations. The error function (12) does not converge as *l′*_*T*1_ decreases. (B) For small values of the T1 transition threshold, some simulations fail to complete (see main text). (C) The dependence of the number of cell rearrangements on *l′*_*T*1_ for 100 model realisations. The number of cell rearrangements is larger than 100 for a large value of the rearrangement threshold, *l′*_*T*1_ *>* 0.1, whereas cell rearrangements are suppressed for small values of the rearrangement threshold, *l′*_*T*1_ < 0.001, with cell rearrangement numbers less than 30. (C) Varying amounts of cell rearrangement lead to different distributions in cell neighbour numbers. Parameter values are listed in table 1. Error bars denote standard deviations across 100 simulations.

A large T1 transition threshold of 0.2 length units leads to a large number of T1 transitions, whereas T1 transitions are suppressed for thresholds of 0.003 length units or smaller (figure 8C). This variation in the number of cell rearrangements influences summary statistics of cell packing, for example leading to variations in the cell neighbour number distribution. For large rearrangement thresholds, e.g. *l′*_T1_ = 0.2, the number of cell rearrangements is high, leading to a high proportion of hexagons (around 0.6), whereas suppression of cell rearrangements for small cell rearrangement thresholds, for example *l′*_T1_ = 0.2, leads to a wider distribution of cell neighbour numbers with a proportion of hexagons below 0.4. The number of cell rearrangements is stable between T1 transition thresholds of 0.02 and 0.003. In this regime, the proportion of hexagons varies slightly between 0.425 and 0.409 (figure 8D). Despite the stable number of T1 transitions across this parameter regime between 0.02 and 0.003 the final vertex positions differ for any two values of the T1 transition threshold, as reflected in values of the error function.

As illustrated in figure 8B, if the T1 transition threshold is smaller than 0.001, simulations fail to complete as the simulation algorithm encounters situations that it cannot resolve, for example due to overlapping or self-intersecting cells. An example of how a simulation can fail due to a small value of the T1 transition threshold is provided in figure 7. A snapshot is taken of the simulation at the last two time steps before simulation failure. Due to a short edge two boundary vertices in the tissue appear merged (arrow in figure 7A). This short edge is magnified for the penultimate (figure 7B) and last time steps (figure 7C) before simulation failure. At this last time step, one of the boundary cells becomes concave. The simulation then fails since our vertex model implementation cannot resolve this configuration. When two boundary cells overlap, the simulation procedure attempts to merge the vertex with its closest cell boundary. This procedure fails because the identified boundary is internal to the tissue rather than a boundary interface.

### Simulation results are robust to variation in length of newly formed edges

When cells exchange neighbours by way of T1 transitions, new edges are formed. Each new edge has length *l′*_new_ = ρ*l′*_T1_. In order to investigate the extent to which changes in the length A of newly formed edges can affect simulation results we define a series of increasing values for *ρ^k^* and the error function

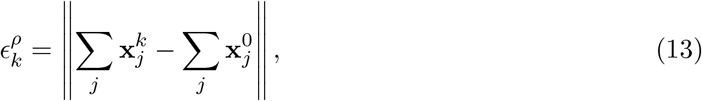

which measures the difference in vertex positions relative to simulations with *ρ*^0^ = 1.05. As shown in figure 9, individual simulations may result in different final tissue configurations than the reference configuration if newly formed edges are twice as long as the rearrangement threshold or longer. Such differences in configuration were observed for three out of 100 simulations, illustrating the robustness of simulation results to the length of newly formed edges.

**Figure 9:**
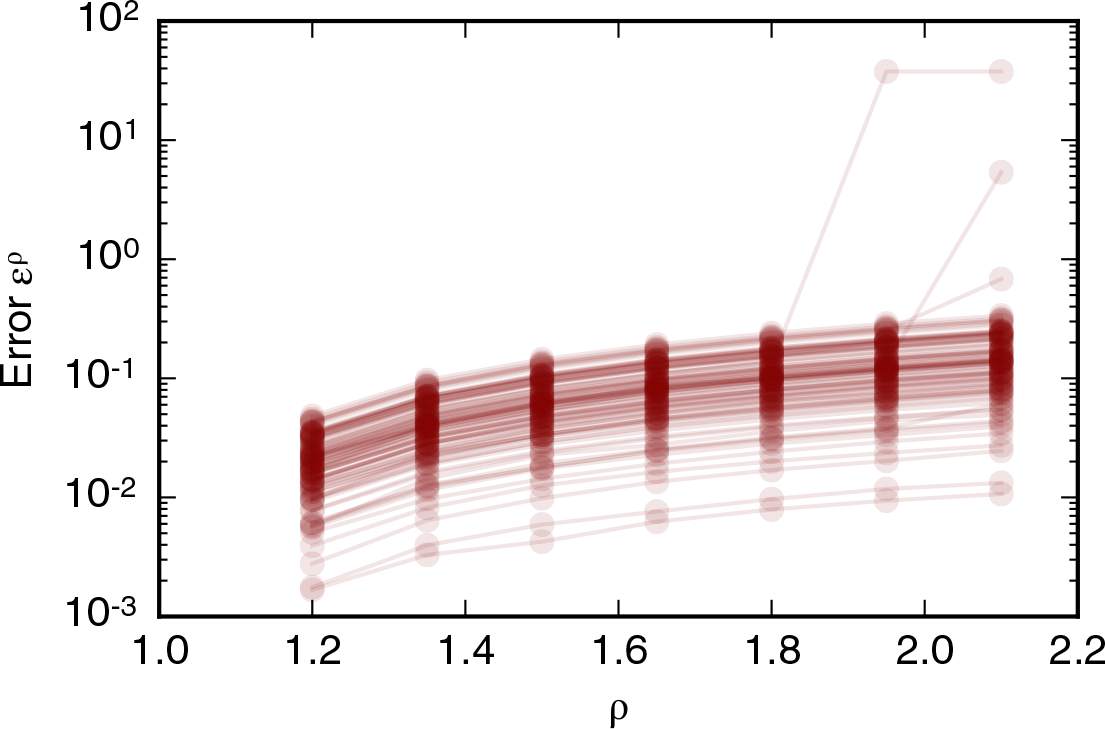
Dependence of simulation results on the length of edges created by T1 transitions, *l′*_new_ = *ρl′*_T_. The error function (13) is recorded for 100 simulations. All simulation parameters are listed in table 1. The error function is smaller than one for *ρ* < 2.0.

### Rate of T2 transitions is robust to variation in the T2 transition threshold over five orders of magnitude

Next, we turn to the value of the T2 transition threshold. We define a series of decreasing values of 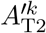 and the error function

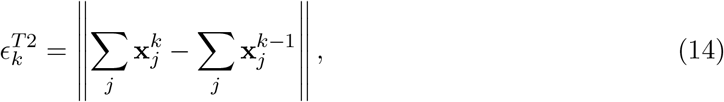

which measures the difference between the final vertex positions of simulations with decreasing values of the T2 transition threshold, 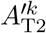. To analyse the value of the error function (14) in a simulation with a significant amount of cell rearrangement and removal we run simulations with *n*_*d*_ = 8 generations, a cell cycle duration of *t′*_*l*_ = 700, and total simulation time *t′*_tot_ = 19600. All other parameter values are listed in table 1.

The value of the error function, on average, is small (figure 10A). However, the error function does not converge for individual simulations and may be large between consecutive values of the threshold. In particular, the error function does not converge to zero. As the threshold decreases, the overall number of T2 transitions in the simulations is stable at approximately 150 T2 transitions per simulation (figure 10B). However, for individual simulations, the total number of T2 transitions may vary by up to 10 as the threshold *A′*_T2_ is decreased. The overall number of T2 transitions does not change over a large range of T2 transition thresholds that covers multiple orders of magnitude, and all simulations complete without errors even if the T2 transition threshold is smaller than 10^−6^, which is three orders of magnitudes smaller than the standard value for this parameter in our simulations. The independence of the number of T2 transitions of the threshold 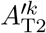 is reflected in tissue-level summary statistics, such as the distributions of cell neighbour numbers, which are unaffected by changes in the T2 transition threshold (figure 10C).

**Figure 10:**
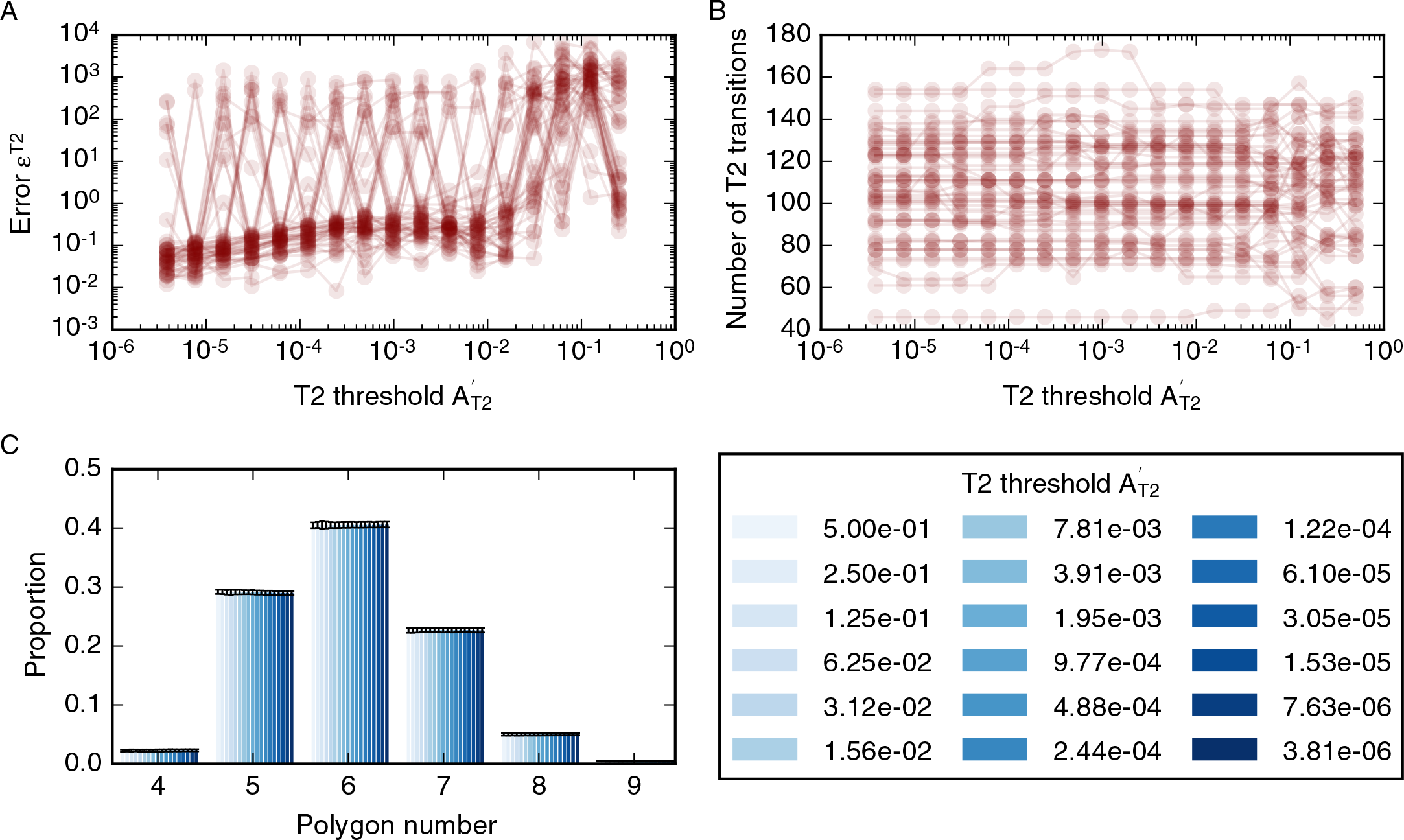
Dependence of simulation results on the T2 transition threshold, *A′*_*T*2_. (A) The dependence of the error function (14) on the T2 transition threshold for 50 model realisations. The error function assumes values less than one for *A*_*T*2_ < 10^−2^ but does not converge. (B) The total number of T2 transitions for 50 model realisations is stable for all observed values of *A*_*T*2_. (C) Tissue-level summary statistics such as the cell neighbour number distribution are not affected by changes in the threshold. Error bars denote standard deviations across 50 simulations. Simulations are run with *n*_*d*_ = 8 rounds of division, a cell cycle duration of *t′*_*l*_ = 700, and total simulation time *t′*_tot_ = 19600. All further simulation parameters are listed in table 1.

### Dependence of the simulation results on the update ordering in each time step

Finally, we investigate whether the update ordering within algorithm 1 may affect simulation results. To this end, we randomise the order in which T1 transitions are conducted during one time step. We find that the update order does not lead to differences in final vertex positions in 100 simulations. This is intuitive, considering that the order in which individual events are conducted is most likely to be relevant in situations where events happen directly adjacent to each other, for example if two adjacent edges undergo T1 transitions at the same time step, if there are two adjacent divisions, or if a dividing cell also participates in cell rearrangement. In these examples, the order in which these events occur during one time step may have an impact on simulation outcomes. Our results imply that no adjacent two edges undergo T1 transitions in 100 sample simulations.

## 4 Discussion

Cell-based models have the potential to help unravel fundamental biophysical mechanisms underlying the growth and dynamics of biological tissues. However, the numerical implementation of such models is rarely analysed and the dependence of model predictions on implementation details often remains unexplored. Here, we analyse a widely applied class of cell-based models, a vertex model, and probe to what extent experimentally relevant summary statistics can depend on implementation details, such as the choice of numerical or non-physical model parameters.

For example, we find that the speed at which cells grow and divide relative to the speed of tissue relaxation can significantly alter *in silico* tissue behaviour. The total number cells in the tissue, as well as the tissue area and the number of cell rearrangements, varies by up to a factor of two as the mean cell cycle duration is changed. Summary statistics of cell packing, such as the distribution of cell neighbour numbers, or the correlation between cell neighbour number and area, are less strongly affected by the exact choice of timescale; the main features B of these statistics are preserved in all cases. This finding that the total cell number and tissue area depend on the mean cell cycle duration suggests that cell extrusion may be induced in fast-growing tissues.

The distribution of cell numbers for the case of quasistatic simulations, identified as simulations where increases in the cell cycle duration would not lead to an overall increase in tissue area or cell number, differs from previously reported results [3]. Specifically, we observe fewer pentagons than hexagons. This discrepancy might arise from a difference in how equation (2) is used to evolve the tissue. For example, our implementation of the cell cycle differs from other implementations where the cell cycle duration varies spatially in the tissue [4, 24, 28]. Further, in [3], a global energy minimisation scheme is used to propagate vertex positions, whereas a more accurate force-based approach is used here. A major difference between the two approaches is the fraction of cells in the tissue that are allowed to grow and divide concurrently. In our implementation, up to one third of the cells undergo cell-growth at any given time, whereas in other implementations all cells grow and divide sequentially. Further analysis is required to understand to what extent synchronous growth and division can affect cell packing in epithelial tissues. Milan et al. report that up to 1.7% of cells in the early wing disc are mitotic at any given time [57]. However, mitosis and cell growth may not happen consecutively, hence the optimal choice of the duration of the growth phase in our simulations is unclear. Overall, it is unclear to what extent different choices for the cell cycle model may influence summary statistics of cell packing.

Our analysis of forces throughout simulations, presented in figure 3, reveals that, on average, the area force contribution is stronger than the edge force contribution and the perimeter force contribution on a given vertex. Further, forces on cells increase during phases of proliferation and growth. Our findings may be of relevance in force-inference approaches that estimate forces using segmented microscopy images of epithelial tissues [58–60]. Force-inference methods often assume that the measured configuration of cells is in equilibrium and it is unclear to what extent force-inference approaches introduce errors if this is not the case. In our simulations, forces are up to 50% higher when simulations are run in a dynamic regime, where cells grow and divide, than in the static regime at the end of the simulation, where cells are relaxed into a static configuration.

The vertex positions, as well as simulation summary statistics, vary as the time step is changed, and differences in vertex positions decrease with the time step. Counterintuitively, large time steps can suppress cell rearrangement in vertex simulations. This may be explained by considering that, for large time steps, vertex positions move further than the length threshold for cell rearrangements, and instances when the lengths of cell-cell interfaces fall below this threshold may not be resolved. Importantly, in order for differences in simulation results to be negligibly small, a time step has to be chosen that is five orders of magnitude smaller than the average cell cycle duration in our simulation, and six orders of magnitude smaller than the simulation time. For individual simulations, simulation outcomes may change if a smaller time step is chosen, an effect that is preserved even when a higher-order numerical scheme, such as fourth-order Runge-Kutta, is used. The latter finding confirms that, for vertex model implementations with ad-hoc rules for cell rearrangement and division, such as in this study, the benefits of higher-order numerical schemes diminish, and it is beneficial to reduce the computational cost of the algorithm by using a simpler numerical scheme, such as forward Euler. A forward Euler scheme is more computationally efficient than a fourth-order RungeKutta scheme since it requires fewer floating point operations per time step. In our simulations, differences in simulation outcomes with decreasing time steps occurred at all observed choices of the time step for both numerical schemes investigated. More research is required to analyse the extent to which further decreases in the time step can lead to convergence of the simulation results. Here, we stopped investigating the effects of further decreasing the time step due to prohibitive increases in calculation times as the time step is decreased. In previous studies, vertex models have been reported to converge as the time step is decreased [45, 56]. Our analysis differs from these previous studies by considering a tissue undergoing cell division and rearrangement rather than relaxation from an initial condition.

The simulation results are sensitive to the T1 transition threshold chosen in the simulation. The size of the T1 transition threshold can be used to regulate the extent to which the simulated tissue is allowed to rearrange in order to minimise energy. Literature values for this quantity span a range from 0.1 [4,48] to 0.01 [31]. Final vertex positions of individual simulations change with the value for the T1 transition threshold and do not converge as the threshold is decreased.

Our results that both the time step and the cell rearrangement threshold may influence the rate of T1 transitions illustrates that these parameters are interconnected. When the time step is chosen sufficiently large such that vertices move further than the cell rearrangement threshold between time steps, cell rearrangement is suppressed. This means that if a small cell rearrangement threshold is chosen, a sufficiently small time step needs to be chosen. A careful choice of time steps and cell rearrangement threshold is crucial since an incorrect choice may lead to failure of the simulation algorithm. For vertex models designed to simulate polycrystalline materials an adaptive time-stepping scheme has been developed that resolves the exact time at which the end points of a short edge meet, and a T1 transition is performed whenever this happens [18]. More work is required to understand how rates of T1 transitions differ if different conditions for rearrangement are implemented, such as the shortening of an edge to a given threshold or the shrinking edge of an edge to a point. Ultimately, the optimal algorithm to simulate cell rearrangement in epithelial tissues can only be chosen through comparison with experimental results.

While simulated vertex model configurations are sensitive to the size of the time step and thresholds for cell rearrangement, they are less sensitive to the length of newly formed edges, and to thresholds for cell removal. We find that the length of newly formed edges may be up to twice as long as the threshold for T1 transitions without affecting final vertex configurations. However, this may change in other parameter regimes, for example if larger values for the cell rearrangement threshold are chosen.

The size of the area threshold for cell removal may be varied over six orders of magnitude without impacting tissue-level summary statistics, even though the exact number of T2 transitions may differ for any two values of the area threshold. In particular, it seems to be possible to choose arbitrarily small values for the T2 transition threshold without causing the algorithm to fail. There are three effects that may contribute to the stability of small elements in our simulations. First, since small cells with areas close to the threshold for cell removal are far away from their preferred area in our simulations (*A*_0,*α*_ > 1.0), their area force is larger than that of adjacent neighbours. This makes the cells stiff and prevents them from becoming inverted or otherwise misshapen. Second, the relationship between area and cell neighbour numbers presented in figure 2 shows that small elements are most likely to be triangular. Our simulation algorithm does not permit T1 transitions if the short edge is part of a triangular cell in order to prevent triangular elements from becoming inverted and thus the algorithm from failure. Third, this relationship between cell area and cell neighbour number may also contribute to the stability of the algorithm when the area threshold is large, for example 0.2. In this case, individual cells may be smaller than the area threshold without undergoing T2 transitions if they are not triangular.

The energy equation (2) provides a geometrical hyphothesis for the removal of cells from epithelia, in which cells are removed from the tissue if this is energetically favourable. Mechanical effects of cell death are an area of increasing biophysical interest [61], and it is the subject of future work to design vertex models that allow alternative hypotheses for cell death to be tested.

Here, we analysed how numerical and non-physical parameters can influence experimentally measurable summary statistics in cell-based models by examining a force-propagation-based implementation of vertex models. Individual results may be relevant to other implementation choices. For example, our finding that the duration of the cell cycle in our model influences simulation outcomes may mean that parameters that control the rate of energy-minimisation may influence results in other vertex model implementations [3,25,62]. In general, further work is required to understand how other choices of implementation schemes may impact computational model predictions. For example, the noise strength in a Monte Carlo vertex propagation scheme [39, 40] or the choice of energy-minimisation algorithm may influence vertex model behaviour.

While most of our findings are of a numerical nature, some have explicit biological relevance. Our analysis of the dependence of tissue properties and forces on the mean cell cycle duration reveals that the vertex model predicts increased forces in tissues undergoing growth and proliferation, and that fast tissue growth may induce cell extrusion. Our findings further suggest that statistics of cell packing may depend on the nature of the cell cycle or the boundary condition of the tissue. Note that findings that do not make explicit biological predictions, such as the robustness of the vertex model to changes in the area threshold for cell removal, or its sensitivity to changes in the length threshold for cell rearrangement, are nonetheless highly relevant, since these findings highlight that choices of model design and implementation have to be carefully considered when applying vertex models quantitatively.

Throughout the manuscript we use non-dimensional parameters that arise when rescaling time and space by the characteristic length and time scales of the model. The use of such rescaled parameters is beneficial in this case since it allows, for example, the comparison of our model parameters to previously used values [3, 4, 28]. Further, we identify reference parameter values for which our simulations are physically reasonable. By providing non-dimensional values for these parameters we facilitate their reuse in other applications where the physical values of the characteristic length or time scales may be different.

## 5 Conclusions

Our results illustrate that care needs to be taken when drawing predictions using cell-based computational models because implementation details such as the size of the time step or non-physical parameters, such as length thresholds for cell rearrangement, may influence model predictions significantly. With the rise of quantitative analysis and quantitative model-data comparison in biophysical applications, choices of model implementation become increasingly relevant. To enable the use of cell-based models in quantitative settings, it is important to be aware of any influences that implementation choices may have on model predictions when analysing a specific biophysical phenomenon. Understanding model behaviour in detail is crucial to prevent modelling artefacts from influencing experimental predictions and clouding our biophysical understanding and, as such, our findings emphasise the need to fully document algorithms for simulating cell-based models. Close attention to implementation details is required in order to unravel the full predictive power of cell-based models.

## Acknowledgements

J.K. acknowledges funding from the Engineering and Physical Sciences Research Council through a studentship. The authors would like to acknowledge the use of the University of Oxford Advanced Research Computing (ARC) facility in carrying out this work.

